# Protein tertiary packing is nearly achiral

**DOI:** 10.64898/2026.07.13.737334

**Authors:** Sagar D. Khare

## Abstract

The chirality of proteins originates at the Cα stereocentre of each L-amino acid, and is expressed in secondary structure as the consistent twist of α-helices and β-sheets. How far does this handedness propagate as secondary-structure elements pack into a tertiary fold? This question is central to the development of neural networks for the design of heterochiral and mixed-chirality proteins containing mirror-image D-amino acids. Such networks are most effectively trained on the far more abundant structural data available for natural all-L proteins, yet in practice they must generate and evaluate the reflected, D-configured counterparts, which are not explicitly represented in the training set. Here, I develop a spatial tessellation-based parity-odd descriptor, the signed volume of Delaunay tetrahedra (*V_D_*), to measure how chirality is distributed in protein structure. Reflecting a structure only flips the sign of any parity-odd descriptor, so the asymmetry of the *V_D_* distribution quantifies the chirality of a given sequence-local or tertiary structural element. I find that the signed Delaunay volume distribution within individual secondary-structure elements is strongly asymmetric, but this handedness largely cancels once elements pack against one another. The resulting tertiary distribution is nearly symmetric across a broad range of structures, from monomers to protein-protein and protein-ligand complexes. Individual folds can nonetheless be strongly handed at the tertiary scale, solenoid and repeat proteins most of all, yet across a broad sample of the fold universe this handedness cancels, leaving the ensemble near-achiral. Tertiary packing is therefore only weakly chiral, with the small residual handedness greatest at binding interfaces and near-zero in the buried core. How much chirality a model perceives in a D-protein is thus largely a choice of representation, a trade-off between reflection symmetry and the richness of structural information the representation retains. Because *V_D_*reduces the parity-odd content of each structural element to a single scalar while capturing tertiary protein packing, it offers a natural representation for the design of heterochiral complexes and mixed-chirality proteins.

**Significance Statement:** The macromolecules of life are handed, and this mirror-asymmetry is built into every protein at the scale of its amino-acid monomers. Local segments of proteins are also strongly handed. Alpha helices and beta sheets are consistently right-handed. Does this handedness build up through the layers of a protein’s structure, as in some synthetic polymers, or cancel? Using a geometric test across thousands of structures, we find that it largely cancels: the 3D-packing of a folded protein, when described as a tetrahedral tessellation, is almost symmetric under reflection, though its building blocks are not. This means that the design of mirror-image D-protein binders, which are attractive as protease-resistant, low-immunogenicity drug candidates, could rely on AI models trained only on natural L-proteins, using representations developed here.

## Introduction

Natural proteins are built almost exclusively from L-amino acids, and this fixed stereochemistry at the Cα centres makes secondary structures handed: alpha-helices are right-handed and beta-sheets carry a consistent right-handed twist (1), and beta-alpha-beta crossover connections are near-universally right-handed (2, 3). Mirror-image proteins, assembled entirely from D-amino acids, fold to the exact reflection of their L-counterparts (4) and can form specific heterochiral complexes with natural L-targets. Such D-proteins are of growing therapeutic interest because they resist proteolytic degradation and tend to evade immune recognition (5). Progress in designing D-proteins and heterochiral binders has been enabled by steady progress in the chemical synthesis of D-proteins and peptides (6), e.g., an enzyme with reciprocal chiral specificity (7), the development of technology for selection of protease-resistant D-peptide ligands by mirror-image phage display (8), development of potent D-peptide inhibitors (9), the synthesis and folding of a full mirror-image enzyme (10), and the structural validation of a heterochiral D-protein—L-target (VEGF) complex by racemic crystallography (11). More recently, deep-learning design has produced *de novo* heterochiral binders with nanomolar affinity and crystallographically verified accuracy (12).

The dominant computational route for designing a D-protein binder is the mirror trick: reflect the L-target into a D-protein, design an ordinary L-binder against it with standard tools, and reflect the resulting binder back to obtain its D-form. This route is exact in principle, but every deep-learning tool used along the way, whether a structure predictor (13–15) or a sequence- and structure-design network (16, 17), is trained almost exclusively on natural L-structures, so the reflected D-target is an out-of-distribution input. Consequently, structure predictors show high chirality-violation rates on heterochiral complexes even when D-stereochemistry is specified explicitly (18). Recent generative approaches to D-peptide design therefore either wrap an L-trained model in an inference-time reflection (D-Flow; (19)) or add explicit chirality-aware axial features (PepMirror; (20)). How far out of the learned distribution a reflected target lies, however, depends on where chirality resides in a protein structure and how far it propagates from the backbone into tertiary packing, and this has not been quantified.

Molecular handedness of building blocks can propagate to much larger scales in polymers. In cooperative helical polymers such as polyisocyanates, a small chiral bias is amplified along a chemically uniform backbone into a single global helical sense, in the so-called sergeants-and-soldiers mechanism (21, 22). Whether the local handedness of protein secondary structure is likewise amplified as elements pack co-operatively into a fold, or instead cancels, is not known a priori, and motivates the quantification of chirality in tertiary packing.

Chirality can be detected only by parity-odd quantities, those whose sign changes under reflection. Parity-even quantities such as distances and angles are blind to it. Any scalar measure of chirality is a pseudoscalar, and the simplest one that can be formed from coordinates is the signed volume of a tetrahedron (23). To enumerate four-residue neighborhoods across a structure without imposing an arbitrary distance cutoff, I use Delaunay tessellation (**Fig. 1a**). Voronoi and Delaunay tessellations have long been used to analyze protein packing and residue volumes (24, 25), and the Delaunay tessellation of residue positions was introduced for proteins by Tropsha and colleagues (26) and developed into four-body statistical potentials that discriminate native from non-native conformations more accurately than pairwise potentials (27). Those potentials classify each tetrahedron by its residue composition and a sequence-continuity class, both of which are parity-even and carry no handedness. Here, I compute the parity-odd signed volume of the simplex (**Fig. 1b**), and use its distribution in datasets of protein structures to ask (a) how far chirality reaches into tertiary packing by comparing the signed volume distribution of sequence-local and sequence-distal (i.e., tertiary) tetrahedra (**Fig. 1c**), and (b) how much of the chirality that an L-trained model would perceive in a D-protein can be removed by the choice of representation, which trades reflection symmetry against the structural information the representation retains.

**Figure 1.**
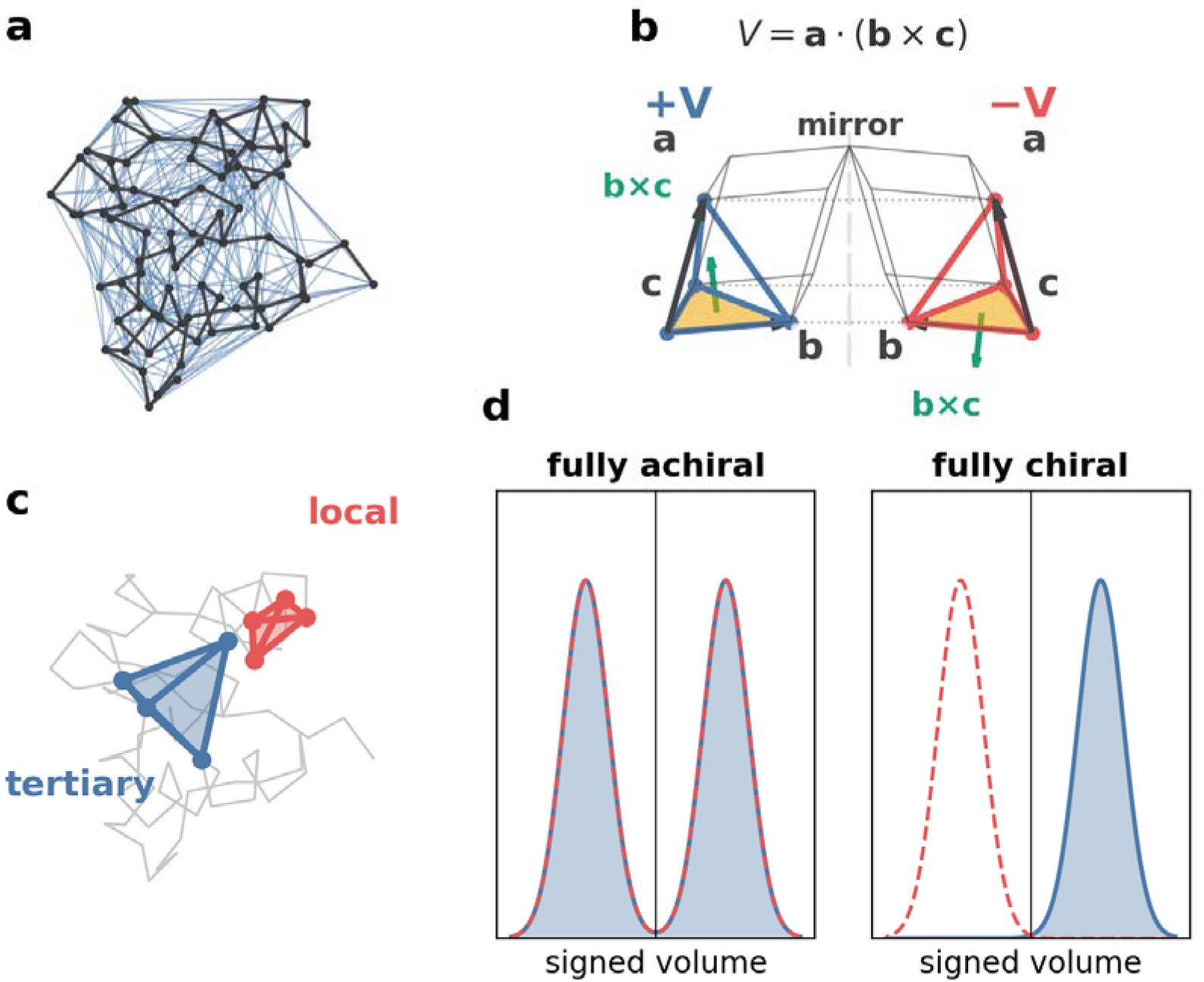
The signed-volume chirality measure. (a) A protein (MDM2, 1YCR) as its Cα trace with the Delaunay tessellation overlaid (short edges only), partitioning space into tetrahedra without a distance cutoff. (b) The signed volume of one tetrahedron, inscribed in the parallelepiped of its three edge vector *a*, *b*, *c*, beside its reflection; reflection flips the sign of *V* while leaving all distances unchanged, so *V* i parity-odd. The sign requires an ordering of the four vertices, taken here by sequence position. (c) Sequence span distinguishes a local tetrahedron of four sequence-adjacent residues (red) from a tertiar tetrahedron of four residues close in space but far in sequence (blue). (d) For a dataset, the signed-volume distribution (blue) is compared with the mirror distribution the reflected D-set would produce (red dashed).

## Results

### Is a mirror-image protein in distribution? A parity-symmetry test

A reflection is an improper isometry: it preserves every interatomic distance while inverting handedness. Any scalar descriptor, *d*, that is parity-odd flips under reflection and therefore takes the value −*d* for the reflected structure. It follows that the set of descriptor values for a collection of D-proteins is the exact sign-flip of the set for the corresponding L-proteins. Therefore, asking whether D-proteins are out of distribution for a given representation is equivalent to asking whether the distribution of a parity-odd descriptor over L-proteins is symmetric about zero (**Fig. 1**). If the distribution is symmetric, every value a D-protein would contribute already occurs among L-proteins and the D-protein is in distribution; if it is one-sided, the reflected values fall where no L-protein value lies. Importantly, no D-structures are required to make this comparison.

To quantify symmetry, I compare a descriptor histogram with its mirror image, the histogram of the reflected, sign-flipped descriptor values. The symmetry is one minus the total-variation distance between the histogram and its mirror,

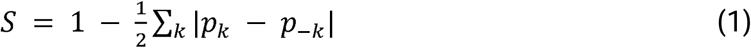

where p_k_ is the fraction of descriptor values in bin k and p_−k_ the fraction in the mirror bin, so that *S* equals 1 for a perfectly symmetric, achiral distribution and 0 for a completely one-sided, fully chiral one. Alongside *S* I report a signed chirality index, *C*, that reads only the overall sign of each signed volume,

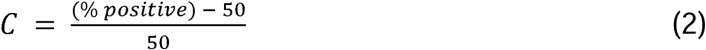

for which 0 is achiral and plus or minus 1 fully one-handed. *C* gives the direction and strength of any handedness, whereas *S* measures the symmetry of the full magnitude distribution.

### A signed-volume measure of local chirality, applied through Delaunay tessellation

Chirality can be detected only by a parity-odd quantity, *i.e.*, one whose sign flips under reflection. A simple representation computable from raw coordinates is a pseudoscalar (23): the signed triple product of three edge vectors, equivalently the signed volume of a tetrahedron, which inverts to its negative under reflection (**Fig. 1b**),

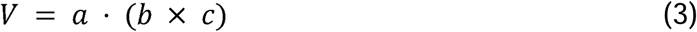

where the cross product ***b*** × ***c*** gives the base-parallelogram area and fixes the orientation while the dot product with ***a*** gives the signed height, so that a tetrahedron is one-sixth of the parallelepiped spanned by ***a***, ***b*** and ***c***. Reflecting the structure leaves every interatomic distance unchanged but inverts *V* to −*V*. A complementary representation, used to make neural network models chirality-aware, is the axial vector or pseudovector such as that cross product, shown as the base normal ***b*** × ***c*** in **Fig. 1b**, which transforms with an extra sign relative to an ordinary polar vector and can be injected as a feature so that a network distinguishes a structure from its mirror (20).

To enumerate four-residue neighborhoods across a structure without an arbitrary distance cutoff, I use Delaunay tessellation, which partitions space into space-filling simplices, tetrahedra in three dimensions, by tessellating Cα positions (**Fig. 1a**) (26). The four-body statistical potentials built on this tessellation classify each tetrahedron by its residue-type composition and a sequence-continuity class (27), both of which are parity-even and so carry no chirality information; the signed volume is the parity-odd complement I use. I normalize it by the cube of the mean tetrahedron edge length to a dimensionless index,

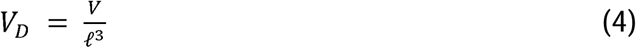

where ℓ is the mean of the six edge lengths of the tetrahedron, so that *V_D_* depends only on the shape of the four points. Its magnitude measures compactness, running from 0 for four coplanar points to a maximum of 1/(6√2) ≈ 0.118 for a regular tetrahedron, the most compact arrangement. Its sign is the handedness of the four vertices taken in sequence order, and flips under reflection. A regular tetrahedron thus reads *V_D_* = +0.118 or −0.118 according to its handedness, and this maximal magnitude fixes the scale of the index. Simplices with any edge longer than 12 angstroms are discarded as convex-hull artifacts.

The four vertices of a tetrahedron can be close together in sequence or far apart, such that a tetrahedron of sequence-adjacent residues is backbone-connected, whereas one whose residues are close in space but distant in sequence is a tertiary contact. I quantify this distinction by the sequence span, the difference between the largest and smallest residue index among the four vertices (**Fig. 1c**). For every tetrahedron composition (e.g. Ala-Ile-Leu-Val) the signed volume *V_D_* of all instances is computed, and its distribution is compared with its mirror image: the two coincide for achiral packing and separate for chiral packing (**Fig. 1d**). Throughout, ‘tertiary packing’ (equivalently, ‘tetrahedral tertiary packing’) is shorthand for this Delaunay four-body measure, the signed volume distribution of tertiary tetrahedra.

To investigate the distribution of chirality in protein structure, I evaluate *V_D_* distributions for four sets of experimentally determined protein structures. (A) The primary set is a sequence-non-redundant collection of 846 heteromeric X-ray complexes (resolution to 2.0 angstroms), each the representative of a distinct 30%-sequence-identity cluster of the ProteinMPNN training data (16); it provides 852,646 interface-proximal tetrahedra. (B) Generalization is assessed on the larger ProteinMPNN training split of 20,383 structures (one representative per training cluster). (C) Ligand-binding-site chirality uses the LigandMPNN small-molecule held-out set of 317 structures (28). (D) As an independent reproducibility control, the principal analyses are repeated on a set of 500 complexes assembled by a different clustering procedure, which shares only 55 of its 500 entries (11%) with the primary set (A).

### Secondary structure carries the chirality, and the V_D_ distribution detects known-handed motifs

There is no a priori reason for tetrahedral tertiary packing to be achiral, because the elements from which a fold is built are themselves strongly handed. For example, an idealized right-handed helix and its left-handed mirror give signed volumes of equal magnitude and opposite sign, confirming that the descriptor reads helix handedness directly (**Fig. 2a**). An idealized twisted beta-sheet, modeled as a saddle-shaped hyperbolic paraboloid following the isotropically stressed description of sheet geometry (29), reads as one-handed and of the opposite sign to the helix. The two peaks in its signed-volume distribution (**Fig. 2a**) are the two geometric families of tetrahedra that the Delaunay tessellation of a near-planar sheet produces, both carrying the same sign.

**Figure 2.**
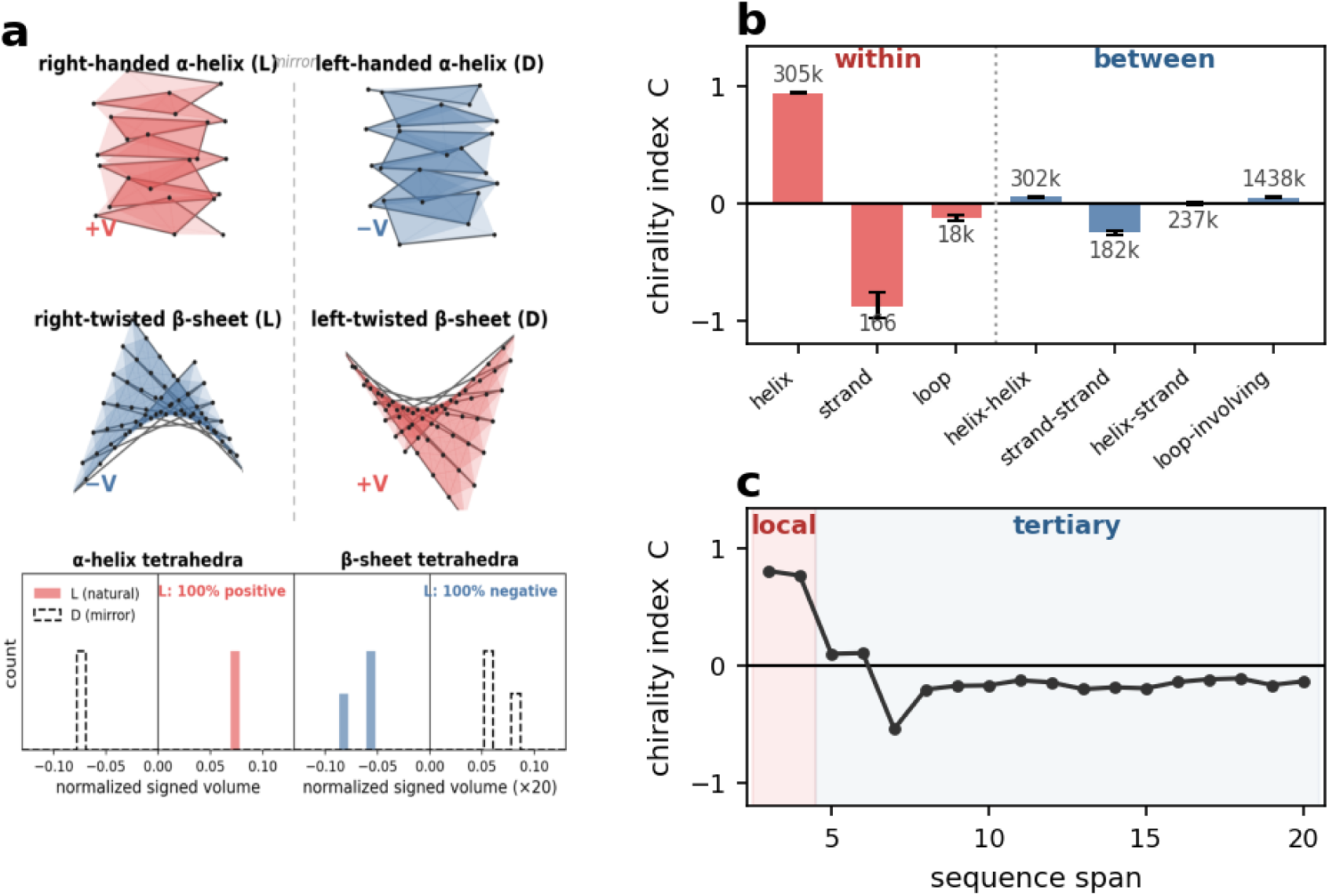
Secondary structure carries the chirality (846 non-redundant structures; *C* = (% positive − 50)/50). (a) Idealized secondary structure and its mirror: right- and left-handed α-helices and right- and left-twisted β-sheets, tessellated and colored by the sign of the local signed volumes, with the normalized signed-volume distributions for the L forms (colored) and their mirror (dashed) below (sheet values scaled ×20). (b) Chirality index within single secondary-structure elements and for tetrahedra bridging two elements. (c) Chirality index resolved by exact sequence span (interface tetrahedra), from the local regime (red) through intermediate span to the long-range tertiary regime (blue).

To examine how the handedness of secondary structures is distributed in experimental protein structures, I assigned secondary structure with a hydrogen-bond (DSSP) analysis and grouped the tetrahedra by secondary structure assignment (**Fig. 2b**), reporting the chirality index *C* defined above. Tetrahedra lying within a single helix are almost entirely of one sign (*C* = +0.94, rising to +0.998 for the sequence-local subset). Tetrahedra within a single strand are strongly handed in the opposite sense (*C* = -0.88), and those within loops weakly so (*C* = -0.12). When tetrahedra instead bridge two different secondary-structure elements, so that handed pieces pack against one another, the handedness largely disappears. Resolving these inter-element tetrahedra by the types of element they connect, strand-strand packing, that is, beta-sheet pairing, retains a clear handedness (*C* = -0.25), recovering the familiar right-handed twist of beta-sheets (1), helix-helix packing is mild (*C* = +0.05), and helix-strand (*C* = 0.00) and loop-involving (*C* = +0.05) packing are achiral (**Fig. 2**). The handedness of a protein is built into its secondary structure but largely cancels, rather than compounding, once those elements pack into a fold.

The same trend in chirality is detected when tetrahedra are classified by sequence span rather than by secondary-structure element (**Fig. 2c**). For a span cutoff between 3 and 8, the sequence-local pool (tetrahedra whose four residues span at most the cutoff) stays strongly one-handed, from 91% of one sign when only span-3 tetrahedra are included to 76% at a span-8 cutoff. The remaining tetrahedra, those spanning more than the cutoff, stay within a few percent of the symmetric value. The local handedness is thus a robust property of backbone-connected geometry. I set the boundary between local and tertiary at a sequence span of four for the rest of the paper: sequence-local tetrahedra have span at most four, tertiary tetrahedra more. Where genuine tertiary handedness does exist the measure still reports it: the beta-alpha-beta crossover, a motif of essentially fixed handedness (2, 3), comes out strongly chiral (98.9% of one sign and near-uniformly right-handed; **Fig. S1**), so the near-symmetry of ordinary tertiary packing reflects the geometry and not an insensitivity of the descriptor.

### Tetrahedral tertiary packing is nearly achiral across environments and functional sites

The tetrahedra of the ProteinMPNN 846-structure set divide sharply (**Table 1**, **Fig. 3a**). Sequence-local tetrahedra, with span at most four, are strongly handed, 89.2% of one sign, and their signed-volume distribution overlaps its mirror poorly (**Table 1**); these backbone-connected neighborhoods carry the handedness of the fold itself. Tertiary tetrahedra, whose residues lie far apart in sequence but close in space, are much closer to symmetric, though not perfectly so, at 47.2% positive, close to but significantly below the achiral value of 50 (**Table 1**). Tetrahedral tertiary packing thus carries a small but statistically significant handedness bias, roughly an order of magnitude weaker than the sequence-local signal and of the opposite sign. Repeating the entire analysis on an independently constructed set of 500 complexes, assembled by a different clustering procedure, gave an equivalent result (**Fig. 3b**; sequence-local 87.2% positive, tertiary 46.1% positive), so the finding does not depend on the particular choice of set.

**Figure 3.**
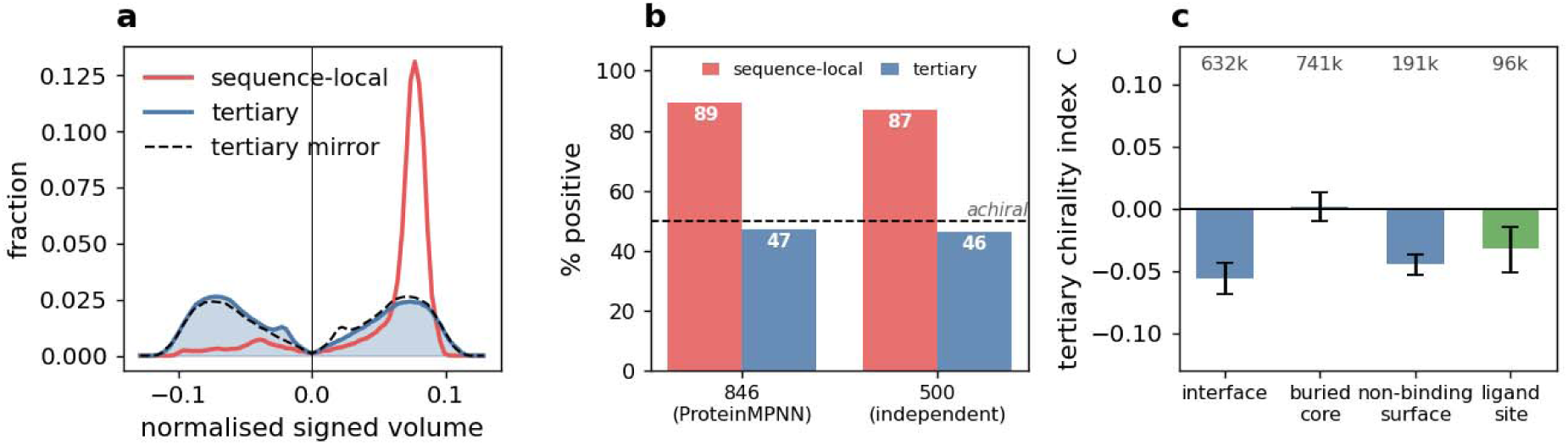
Delaunay packing across protein environments and ligand sites. (a) Normalized signed-volume distributions for sequence-local (red) and tertiary (blue) tetrahedra, the latter with the mirror distribution the reflected D-set would produce (dashed). (b) Sequence-local and tertiary % positive (interface tetrahedra) for the ProteinMPNN 846-complex set and an independent 500-complex set. (c) Tertiary chirality index across the interface, buried core, non-binding surface, and ligand-binding sites (per-bar tetrahedron counts shown).

**Table 1.**
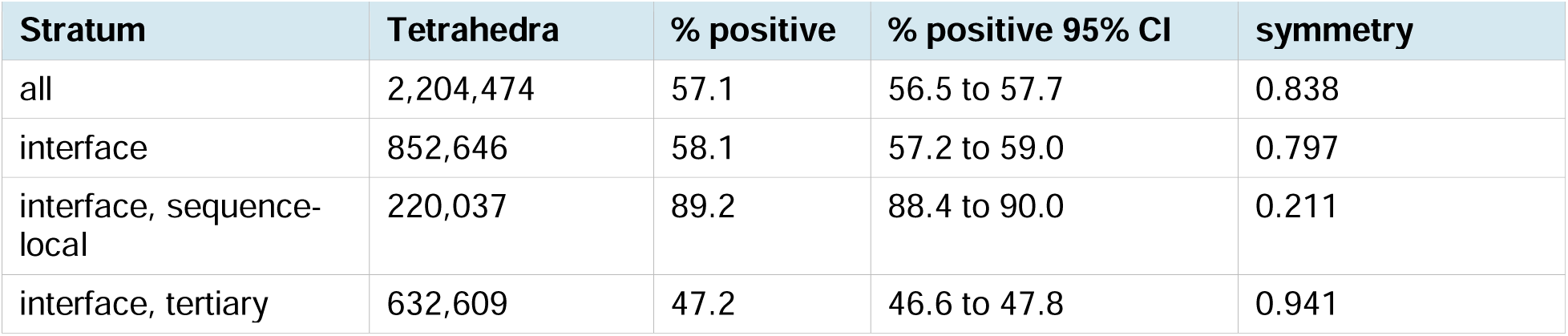
Delaunay signed-volume chirality across a non-redundant set of 846 heteromeric L-L complexes (Cα sites; ProteinMPNN training clusters, X-ray <= 2.0 angstrom, 30% identity). A positive fraction of 50% indicates a symmetric, achiral distribution; confidence intervals are structure-level bootstrap; symmetry is one minus the total-variation distance between the signed-volume histogram and its mirror (1 = perfectly symmetric). The pooled ‘all’ and ‘interface’ rows inherit the strong sequence-local handedness and are interpretable only through the span-resolved split (sequence-local versus tertiary).

This near-achirality of tetrahedral tertiary packing holds across structural contexts, but the small residual handedness is not distributed uniformly within the fold. When each residue is classified by burial and by interface proximity, sequence-local geometry remains strongly handed in every environment and is strongest in the buried core, as expected from its high secondary-structure content (**Fig. 3c**). Tertiary geometry, by contrast, is near-symmetric everywhere, yet the residual that remains is structured rather than random: it is largest at protein-protein interfaces (chirality index *C* = -0.056) and on the non-binding surface (*C* = -0.045), and falls essentially to zero in the buried core (*C* = +0.002), which is the most achiral environment despite being the most densely packed (**Fig. 3c**). The handedness of tetrahedral tertiary packing is therefore concentrated where the backbone is most exposed and participates most directly in inter-chain contacts, and it vanishes in the side-chain-dominated interior.

This pattern extends to functionally specialized surfaces. On a set of 317 small-molecule complexes, tertiary geometry at ligand-binding sites, defined as residues within 5 angstroms of a drug-like ligand, is only weakly handed (*C* = -0.032), comparable to the non-binding surface and far weaker than the sequence-local signal. A drug-binding pocket therefore carries essentially the same chirality signature as bulk tetrahedral tertiary packing, and the ligand-binding geometry of a reflected D-protein is in distribution to nearly the same degree as the rest of the structure.

### Near-achirality of tetrahedral tertiary packing is robust to the choice of representation and residue ordering

A reflected D-protein is in distribution when the signed-volume distribution is symmetric (**Fig. 1d**). However, the sign of *V_D_* is not a property of the four points alone. It combines the handedness of the shape they form with the parity of the rule used to order them. A symmetric distribution of signed volumes can therefore arise in two ways. First, the shapes may be balanced in handedness, a shape and its mirror occurring equally often, which is true achirality. Second, the shapes may be handed while the ordering is uncorrelated with that handedness, scrambling the sign to fifty-fifty, which is an artifact of labeling. The second is highly likely for sequence orderings of long-range contacts, where the packing residues bear no fixed relation to the geometric hand.

To delineate the effects of ordering and shape, I order the four vertices of each tetrahedron by two sequence-independent geometric rules: by distance to the tetrahedron centroid, and by summed distance to the other three, each breaking ties with the other (**Fig. 4a**). Both labelings come only from the pairwise distances within the tetrahedron, so they are reflection-equivariant. Under these rules the sign is strictly a function of the shape, not the order of residues in the sequence to obtain vertex labels. Tertiary tetrahedra still come out near balance (48.8% and 47.8% positive for the centroid and summed-distance rules, respectively, against 48.6% for sequence order, over the 1,751,041 tertiary tetrahedra of the ProteinMPNN-846 set; **Fig. 4b**, **Fig. S4**). The same rules keep the sequence-local fragments strongly one-sided (**Fig. 4b**). Thus, local four-point shapes are handed but tertiary ones are not. A third ordering, by residue burial, is blind to the shape of both local and tertiary tetrahedra. Tetrahedral chirality distributions, local and tertiary alike, using this ordering are fully symmetric (**Fig. 4b**), indicating that the observed chirality differences between local and tertiary tetrahedra are not labeling artifacts.

**Figure 4.**
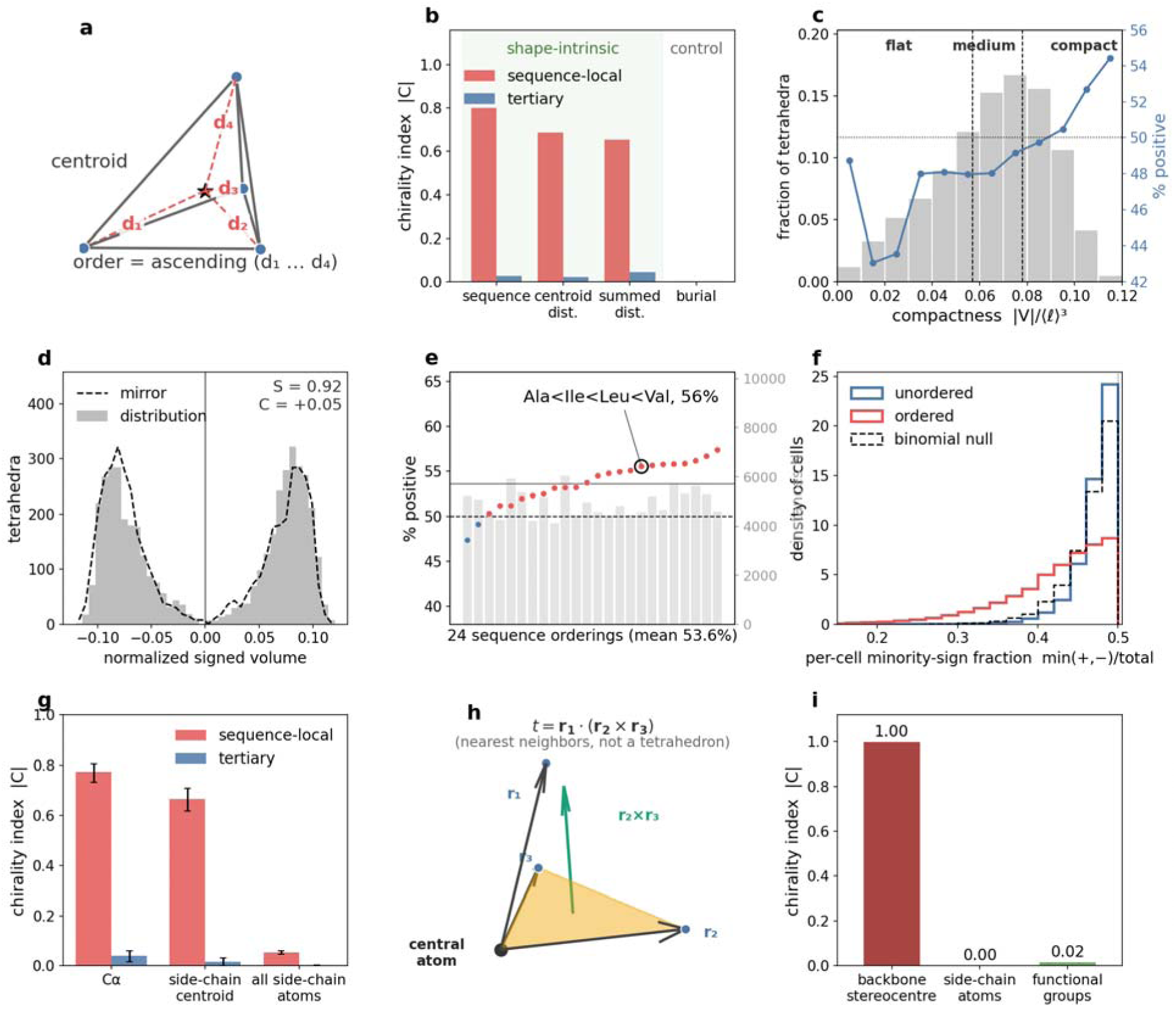
Tertiary packing is robustly near-achiral and the residual is backbone-local. (a) Shape-based vertex ordering by internal distances (distance to centroid, *d*_1_-*d*_4_). (b) Chirality index |*C*| of sequence-local (red) and tertiary (blue) tetrahedra under four orderings (sequence, centroid distance, summed distance, and a geometry-blind burial control; ProteinMPNN-846 set). (c) Distribution of compactness |*V*|/OℓO³ over tertiary tetrahedra (grey) with tertile cuts (dashed) and % positive across it (blue). (d) One composition (Ala-Ile-Leu-Val, 846 set): normalized signed volumes with their mirror (dashed). (e) The 24 sequence orderings of Ala-Ile-Leu-Val (long-range, ∼20K set), instance count *N*, canonical ordering circled. (f) Per-cell minority-sign fraction min(+,−)/total over all compositions for unordered (blue) and ordered (red) composition, with a matched binomial null (dashed). (g) |*C*| on Cα, side-chain centroids, and all side-chain heavy atoms (bootstrap 95% intervals). (h) The nearest-neighbor signed triple product *t* = *r*1·(*r*2×*r*3) of an atom and its three nearest neighbors. (i) |*C*| of that triple product for the backbone Cα stereocentre, side-chain atoms, and aromatic functional groups.

To investigate the robustness of the observed shape-based near-achirality, I classified all tetrahedral shapes based on their compactness. A nearly coplanar tetrahedron has a vanishing signed volume whose sign could arise from uncertainty in crystallographic structure determination, so the near-achirality might reflect the signal from these flat, sign-ambiguous tetrahedra rather than the compact, well-formed neighborhoods. However, ranking tertiary tetrahedra by compactness rules shows that the compact tetrahedra are the most balanced (50.5% positive; **Fig. 4c**). The small residual handedness in tertiary packing therefore sits in the flat, near-degenerate tetrahedra, not in the compact ones that dominate packing.

Another approach to delineate the confounding impact of sequence-based ordering on the signed-volume distributions is to resolve every ordering of every four-residue composition and ask whether the balanced chirality pattern persists. With a 20-residue alphabet there are 8,855 possible unordered compositions (multisets of four residues) and 20a = 160,000 ordered compositions (a residue at each sequence-ordered position i < j < k < l). Essentially all unordered (8,855) and 159,995 of the ordered compositions are populated in the ∼20,000-structure ProteinMPNN set (**Table S1**). For one randomly chosen example unordered composition, Ala-Ile-Leu-Val, the tertiary signed volumes are symmetric about zero and overlap their mirror (**Fig. 4d**). Within a fixed sequence order, all tetrahedra of this composition share both their composition and their vertex ordering. Chirality values for a given ordering therefore reflect a genuine handedness of that ordered geometry, not a cancellation among one-sided subpopulations due to order scrambling. All 24 orderings stay within a narrow band around 50% (**Fig. 4e**, **Fig. S3**), for example, the Ala < Ile < Leu < Val ordering (circled in **Fig. 4e**) is 56% positive. Across all compositions and orderings, the signed volume distributions remain two-sided at both the tertiary and long-range scales, although fixed ordering introduces increased heterogeneity of handedness compared to the unordered cases (a larger between-cell spread; **Fig. 4f, Fig. S3**). The most handed tetrahedra are also the least compact, so geometry, not composition, sets the small residual (**Fig. 4c**).

A residue can enter the tessellation as its Cα, its side-chain centroid, or through all its side-chain heavy atoms. The first two place one point per residue, so each tetrahedron is a four-residue neighborhood. The third tessellates every side-chain atom, so a tetrahedron is a four-atom simplex that may lie within one residue or span several; each atom keeps its parent residue index for the local-tertiary split. On Cα points the sequence-local tetrahedra are strongly handed. On side-chain centroids the signal weakens. On all side-chain atoms it is near-symmetric even locally, because the backbone, where the local handedness lives, no longer defines the tetrahedra (**Fig. 4g**). Tertiary packing is, however, near-symmetric under all three choices. Removing the backbone from the representation thus removes the handedness.

Handedness is also sharply localized at the atomic scale: the distribution of signed triple products of an atom and its three nearest neighbors (**Fig. 4h**) is fully one-sided at the backbone Cα stereocentre but achiral around side-chain atoms and functional groups (**Fig. 4i**) and the distribution of sidechain atomic triplets is achiral (**Fig S2**).

Thus, across all tests, namely vertex ordering by shape, fixed ordering within an amino acid composition, and choice of the residue central atom, a reflected D-protein is largely in distribution for Delaunay tertiary packing geometry, while the handedness is concentrated in the backbone Cα trace.

### Fold-level handedness cancels across the protein universe

The near-achirality of tertiary packing reflects the distribution of tetrahedral neighborhoods over a diverse collection of protein folds, and it is worth asking what it looks like fold-by-fold. An individual fold can be strongly handed at the tertiary scale, for example, solenoid and repeat proteins whose architecture is set by a single long-range rule that stacks repeated units into a superhelix. Restricting each chain to its CATH domain and tessellating only those Cα sites, the domain-isolated tertiary chirality index reaches |*C*| ≈ 0.2 to 0.4 for these folds (**Fig. 5a-c**), with a sign fixed by the hand of the superhelix. A right-handed parallel β-helix (pectate lyase, PDB 1AIR) and a left-handed parallel β-helix (LpxA, PDB 1QRE), the same architecture built with opposite hand, carry opposite-sign tertiary chirality (*C* = +0.23 and −0.38), and a TPR α-solenoid (PDB 1W3B) reaches +0.39. Fold handedness for individual architectures is therefore substantial, neither absent nor uniformly small.

**Figure 5.**
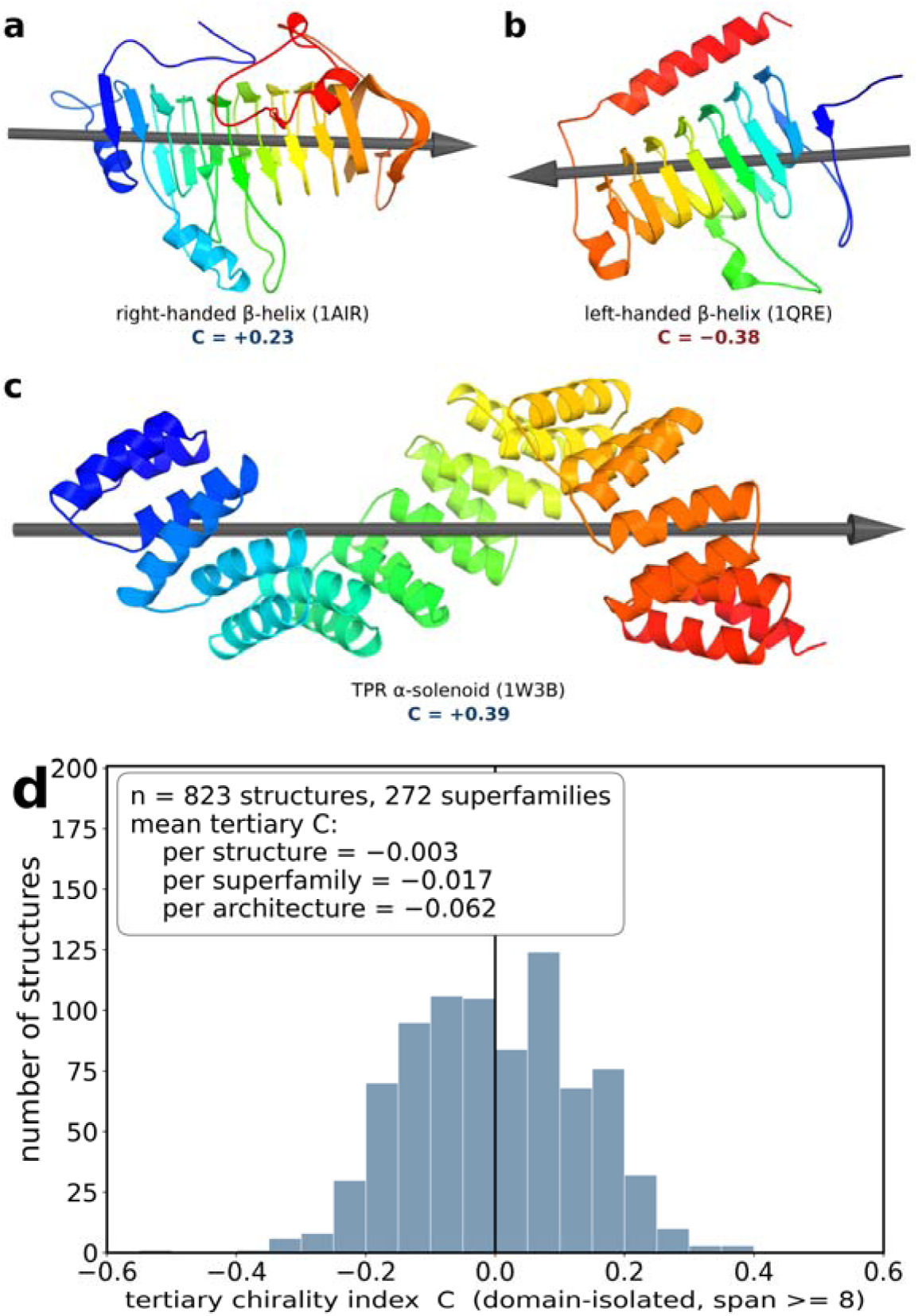
Fold-level tertiary handedness cancels across the protein universe. (a) A right-handed parallel β-helix (pectate lyase, 1AIR; *C* = +0.23) and (b) a left-handed parallel β-helix (LpxA, 1QRE; *C* = −0.38) are the same architecture built with opposite hand and carry opposite-sign tertiary chirality; (c) a TPR α-solenoid (1W3B; *C* = +0.39). Structures are Cα cartoons colored N (blue) to C (red) with the sequence-ordering axis drawn as a grey arrow. (d) Distribution of the domain-isolated tertiary chirality index *C* (tetrahedra of sequence span ≥ 8) over a random sample of X-ray structures (823 scorable structures, 272 CATH superfamilies), each restricted to its largest CATH domain; the distribution is symmetric about zero, with means weighted per structure (*C* = −0.003), per CATH superfamily (*C* = −0.017), and per CATH architecture (*C* = −0.062). All values are measured on sub-structures, each chain restricted to its largest CATH domain, so they report the handedness of an isolated fold or repeat architecture and are larger in magnitude than the tertiary chirality of the intact protein.

Across the fold universe, however, the signs of tetrahedral neighborhoods in these handed folds cancel. A random sample of X-ray structures (823 structures spanning 272 CATH superfamilies), each restricted to its largest CATH domain and scored for tertiary chirality, gives a distribution that is symmetric and centered at zero (**Fig. 5d**; mean per-structure *C* = −0.003). β-sheet folds skew mildly negative and α- and α/β-folds mildly positive, with solenoids in the tails of the distribution, but the ensemble mean is nearly achiral. The small residual depends on how the average is taken, in a way that is itself informative. Weighted by PDB frequency, the mean is zero (*C* = −0.003). Weighting each CATH superfamily equally moves it to *C* = −0.017, and weighting each CATH architecture equally to *C* = −0.062. That drift reflects the larger number of distinct β-sheet fold types in the CATH taxonomy rather than a directional bias in protein structure, and under every weighting the departure from zero is at least an order of magnitude smaller than the handedness of a single solenoid. Near-achirality of tertiary packing is therefore not the absence of handedness in individual folds but its cancellation across the distribution of tetrahedral neighborhoods in a spectrum of individually handed folds.

## Discussion

I develop a measure of packing chirality using spatial tessellation, the Delaunay signed-volume distribution, which partitions a protein structure in a cutoff-free manner into tetrahedra. Across protein structural environments and definitional choices for the tessellation, this analysis finds that the chirality of a protein structure is overwhelmingly a local, backbone-associated property: the chirality signal fades both as residue positions separate in sequence and as the representation moves from the backbone to the side chains. Secondary-structure elements are each strongly handed, but when those elements pack against one another in diverse topologies, the handedness largely cancels, and single atoms, side-chain centroids, functional-group orientations, and long-range tetrahedra are all at or close to symmetric. Tertiary packing is therefore nearly, though not perfectly, achiral.

The small residual handedness that does survive in tertiary packing is not spread evenly. It is concentrated at protein-protein interfaces and on the solvent-exposed surface, while the densely packed core is the most achiral environment of all (**Fig. 3c**). This may seem counterintuitive at first glance, because the core is where secondary structure content and packing is tightest so handed contacts might be expected to accumulate, but it follows naturally if the residual chirality is of backbone origin. In the interior, contacts are dominated by interdigitating side chains whose atomic environments are achiral within error (**Fig. S2**), so the handedness of the backbone is diluted by an essentially symmetric cloud of side-chain atoms. At interfaces and on the surface the backbone is comparatively exposed and takes a more direct part in packing, so the residual handedness is less screened. The interface is therefore the environment in which a reflected D-protein departs most from the natural distribution of L-proteins, although even there the departure is small, roughly 47 percent of one sign against 53 percent of the other, and about an order of magnitude weaker than the sequence-local signal. For mirror-image design this means that the modest chirality cost of working with a reflected target is borne disproportionately at the binding interface, which is precisely the region a binder-design model must read most carefully.

Why should tertiary four-point shapes be achiral when sequence-local ones are strongly handed? The physical interactions that organize tertiary structure are achiral: side-chain shape complementarity and close-packed van der Waals contact are governed by size and shape rather than by any consistent handedness, so tertiary contacts assemble chain segments in relative arrangements that are not fixed by a single chiral convention. This contrasts with the sequence-local scale, whose one-handedness is inherited directly from the fixed L-configuration of the backbone. The same absence of a long-range chiral rule might also explain why sequence order carries no information about tertiary handedness: the sequence-ordered and shape-intrinsic signings agree to within about one percent (**Fig. 4b**).

The lack of chirality propagation in proteins follows from the architecture of cooperative stabilizing interactions, a distinction that is reminiscent of the difference between a ferromagnet and a spin glass. In cooperative helical polymers such as polyisocyanates, coupling along a chemically uniform backbone amplifies a small local bias (the sergeants-and-soldiers effect) into a single global chirality (21, 22). A solenoid is the protein counterpart: one long-range rule winds its repeats into a superhelix, so local chiral tendencies align and add to a net hand, like a ferromagnet, and the tertiary signed volume is one-sided with a sign that follows the superhelical order (**Fig. 5**). A globular fold is closer to a spin glass. Its handed elements pack in varied, aperiodic geometries through many heterogeneous, frustrated contacts, long recognized as the statistical-mechanical basis of cooperative protein folding (30–32), and these present no single chiral convention along which to align, so the four-body signs of tertiary contacts are uncorrelated and average out. Cooperativity by itself therefore does not imply net chirality. Through one kind of coupling it yields aligned, macroscopically handed order, but through many competing couplings it selects a unique native structure while its chiral contributions cancel. The handedness in the latter case is sub-structural: an isolated domain or repeat array can freeze into a definite hand, just as local regions of a spin glass carry specific orderings while the system as a whole does not, and it is aggregation, over the whole chain in many cases and across the fold ensemble overall, that returns the average to near-zero.

How much chirality an L-trained model perceives in a reflected D-protein is not fixed but a property of the representation, which sets a trade-off between reflection symmetry and structural information. Chirality can be suppressed to any degree by discarding information: a representation that exposes backbone stereochemistry, the Cα trace or explicit backbone frames, presents a maximally out-of-distribution target, whereas pairwise distances are perfectly symmetric but fix shape only up to a mirror, and a side-chain point cloud is nearly achiral yet omits the packing topology a signed four-body encoding records. Useful representations lie on the frontier between these extremes, as symmetric as possible while keeping the structural information a model needs. For tertiary packing, the Delaunay four-body descriptor developed here has several favorable properties as a representation for training a model (**Fig. 6**): it records which four residues mutually pack, a higher-order structural signal that pairwise features do not capture, while the one parity-odd quantity it carries, the signed volume, is nearly symmetric for tertiary tetrahedra. Its parity-even content, the pairwise distances, residue composition, and burial, is reflection-invariant, and the residual handedness in the signed volume is roughly an order of magnitude weaker than the sequence-local signal and barely changes with the vertex ordering (**Figs. S4 and S5**), so a reflected D-protein is nearly in distribution under this representation. Much of the gap between L and D performance can thus be closed by the choice of representation alone, before any model is retrained.

**Figure 6.**
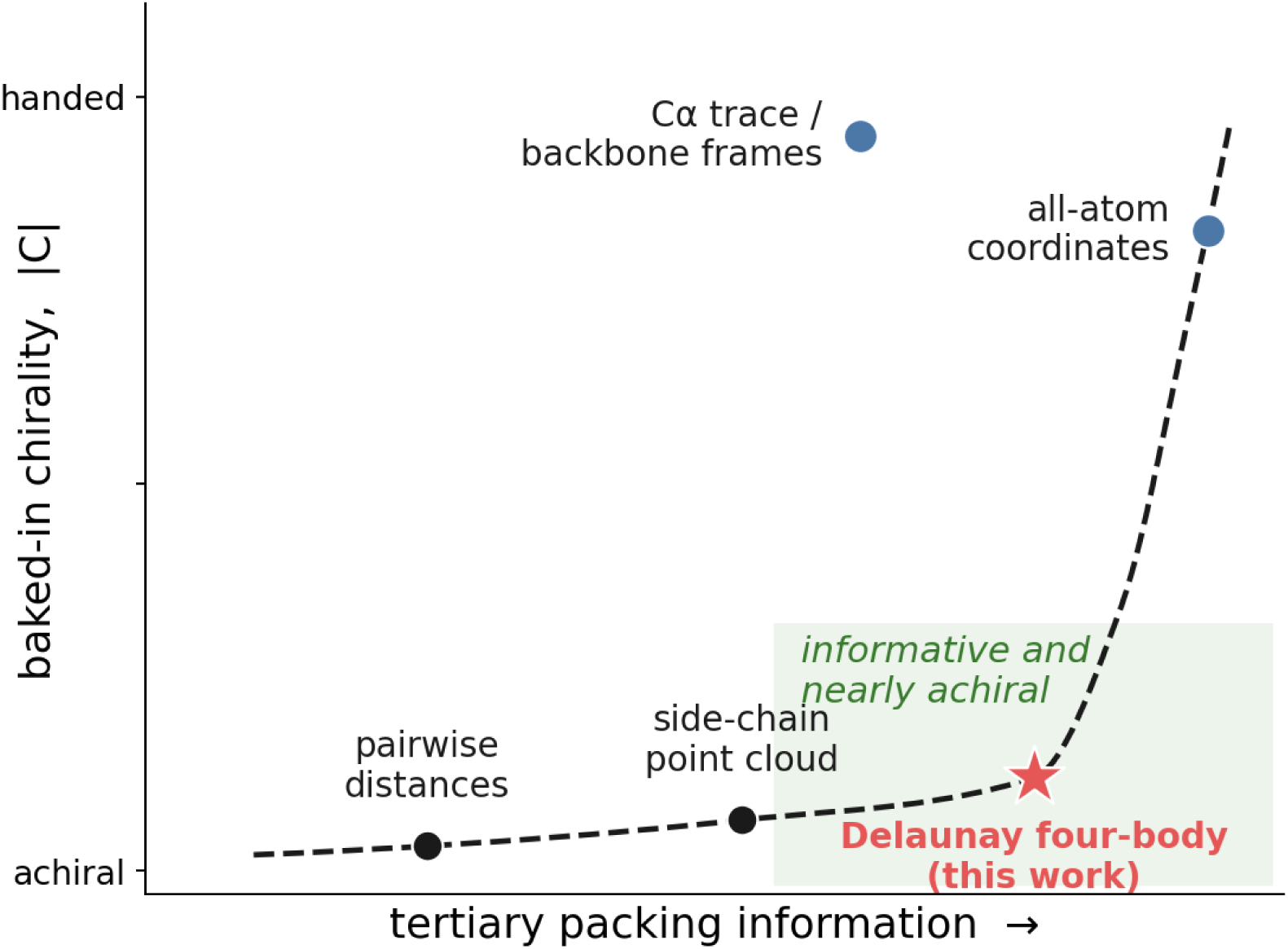
The trade-off between baked-in chirality and structural information across candidate representations of tertiary structure (schematic; the |*C*| axis is anchored by the measured values, the horizontal axis increases with the tertiary structure a representation makes explicit). The dashed curve i the achievable frontier. Tertiary-packing representations (pairwise distances, a side-chain point cloud, and the Delaunay four-body descriptor) lie on the low-chirality part of the frontier; full atomic detail lies at the high-information, high-chirality end; representations that expose backbone stereochemistry (the Cα trace or backbone frames) lie above the frontier.

These hand-designed descriptors point to a learned counterpart: an encoder of tertiary structure that is parity-symmetric by construction, for example E(3)-invariant, or trained with a parity-invariance penalty. The parity-symmetry measure introduced here provides the metric for placing any candidate representation on that frontier, scoring how symmetric it is while its accuracy measures what it retains. The signed volume also complements the four-body statistical potentials built on the same tessellation (27), which encode the parity-even content of each simplex, so the two combine into one chirality-aware four-body encoding. Because these are scalar summaries, their symmetry is necessary but not sufficient for generalization, and the decisive test remains functional: the difference in design quality between L-targets and their D-reflections in an actual pipeline for design of protein-protein complexes, which this analysis lays the groundwork for.

## Materials and Methods

### Structures and datasets

Coordinates were obtained from the RCSB PDB. The primary set is a sequence-non-redundant set of 846 heteromeric complexes derived from the ProteinMPNN training data (16) (the pdb_2021aug02 set clustered at 30% sequence identity): excluding the published validation and test clusters, an entry was taken as heteromeric if its chains span at least two distinct 30% clusters, and a non-redundant set was selected greedily, best-resolution first, keeping an entry only if none of its chains’ clusters had been used and capping resolution at 2.0 angstroms, so no two selected complexes share a 30% cluster. As a reproducibility control, the principal analyses were repeated on a home-built set of 500 entries assembled independently using RCSB server-side clustering (X-ray at resolution at most 2.0 angstroms, at least two protein entities, one representative per 30% cluster, 500 sampled with a fixed seed). Both chains of each complex were used as target in turn; the partner chain defined the interface (target heavy atoms within 5.0 angstroms of any partner heavy atom). ProteinMPNN monomer set contained its training split of 20,383 structures with one representative PDB entry per 30%-identity training cluster, retaining cells with at least twenty tetrahedra. Ligand-binding-site chirality was computed on the LigandMPNN small-molecule held-out set (317 structures).

Idealized secondary structure were built as a standard Cα alpha-helix (radius 2.3 angstroms, rise 1.5 angstroms per residue, 3.6 residues per turn) and a twisted beta-sheet modeled as a saddle surface, the hyperbolic paraboloid z = c x y, with their mirror images obtained by reflection.

### Tessellation and statistics

Secondary structure was assigned by a dependency-free implementation of the DSSP hydrogen-bond algorithm (33) (Kabsch-Sander electrostatic H-bond energy, with helices from i to i+3/4/5 n-turns and strands from the parallel and antiparallel beta-bridge tests), at three-state resolution; elements are maximal contiguous runs of one state, and a tetrahedron is intra-element if all four vertices share one element.

Delaunay tessellation used residue sites (Cα, or side-chain heavy-atom centroid) via the Qhull-backed implementation in SciPy; simplices were labeled interface-proximal if any vertex was an interface residue, and by sequence span.

Confidence intervals in tessellation came from a cluster bootstrap resampling whole structures (3,000 replicates), so that they reflect structure-to-structure variation rather than the pseudo-replication that would result from pooling correlated simplices within a structure as if they were independent.

Environments were assigned per residue by a Cα-neighbor-count burial proxy (core if at least 18 Cα lay within 10 angstroms) and by interface proximity; the local-tertiary boundary was set at sequence span 4 and varied from 3 to 8; the point representation was compared across Cα, side-chain centroid, and all side-chain heavy atoms. Inter-element strata were assigned by tessellating the whole assembly and classifying each tetrahedron by whether its four vertices lie in one secondary-structure element or bridge two, and for bridging tetrahedra by the element types involved (helix-helix, strand- strand, helix-strand, or loop). Beta-alpha-beta crossover handedness was measured for sequential strand-helix-strand element triples (intervening loops permitted) whose strands are parallel (direction dot product above 0.5) and spatially paired (centroids within 12 angstroms), as the signed triple product of the mean strand direction, the inter-strand-centroid vector, and the helix-centroid offset from the strand midpoint.

Ligand-binding-site chirality was computed on the LigandMPNN small-molecule held-out set. A residue was assigned to the binding site if any heavy atom lay within 5 angstroms of a drug-like non-polymer ligand (at least six heavy atoms, excluding water, ions and common buffers and cryoprotectants), and tetrahedra were stratified site / core / non-binding surface as for the protein-interface environments.

### Composition resolved by shape, vertex order, and compactness

Dependence on the vertex ordering was tested by re-signing every tetrahedron under four orderings of its vertices: by sequence position, by distance to the tetrahedron centroid, by summed distance to the other three vertices (each with ties broken by the other distance), and by burial (Cα-neighbor count within 10 angstroms, ties broken by summed distance), the last as a geometry-blind negative control; the per-ordering marginal and the between-composition rate spread are compared in **Fig. S4**. Compactness was the magnitude of the normalized signed volume, |*V*|/mean-edge-cubed, split into three equal-count tertiles at its 33rd and 67th percentiles over all tertiary tetrahedra (0.057 and 0.078), with the positive fraction pooled across compositions within each tertile.

Retaining cells with at least thirty tetrahedra, four quantities were computed. First, marginal handedness: the pooled positive fraction with a 95% confidence interval from a structure-level bootstrap (5,000 replicates) (34). Second, compositional heterogeneity: the null that every cell shares the global positive rate *p*0 was tested with the Pearson chi-square and the likelihood-ratio G-squared (35) on k minus 1 degrees of freedom for k cells, with effect size the method-of-moments estimate of the between-cell standard deviation of the true positive rate, tau, where tau squared is the mean over cells of (p_hat minus *p*0) squared minus *p*0(1 minus *p*0)/n; the fraction of cells more one-sided than 80/20 was compared with a matched cell-size binomial null drawn at *p*0, so the comparison is not confounded by the smaller per-cell counts of a finer partition. Third, whether vertex order adds handedness beyond composition: a nested within-composition G-squared testing, within each unordered composition, whether its sequence orderings depart from the composition’s own rate. Fourth, skew symmetry: a two-sided binomial sign test on the numbers of strongly positive (>70%) and strongly negative (<30%) cells.

## Code and data availability

Every analysis is run by a compute script that takes a structure id list as --idlist and writes a per-structure JSON-lines checkpoint to --out (resumable and time-boxed), followed by a plotting script that writes the figure; the mapping of each display item to its compute scripts, id list, and plotting script is given in **Table S2**. The non-redundant PPI id list (mpnn_ppi_ids.txt) is built by build_mpnn_ppi_idlist.py from the ProteinMPNN metadata; the 20-amino-acid composition list (mpnn_ids.txt) by build_mpnn_idlist.py; and the ligand list (ligand_small_ids.txt) from the LigandMPNN small-molecule test set.

All scripts and plotting code, the identifier lists, per-structure checkpointed results, and a driver that re-fetches the data and regenerates every figure are openly available at Zenodo (doi:10.5281/zenodo.21301663) and https://github.com/khare/protein-tertiary-chirality.

## Notes

https://github.com/khare/protein-tertiary-chirality

https://zenodo.org/records/21301664

### Competing Interest Statement

The authors have declared no competing interest.

