## Supporting Information for "Protein tertiary packing is nearly achiral"

Sagar D. Khare

*Department of Chemistry and Chemical Biology, and Institute for Quantitative Biomedicine, Rutgers University, Piscataway, NJ*

### Contents

**Figure S1.** Supersecondary-motif handedness of known chiral motifs (AlphaFold models)	2

**Figure S2.** Local-shell parity symmetry over the non-redundant set	4

**Figure S3.** Per-composition two-sidedness is robust to vertex ordering, span, and dataset size	5

**Table S1.** Composition-cell counts for the Fig. S3 control	5

**Figure S4.** How the three sign-sensitive vertex orderings differ for tertiary packing	6

**Figure S5.** Tertiary handedness ranked across 30 common protein folds	7

**Table S2.** Reproducibility map	8

#### The beta-alpha-beta crossover: a handed tertiary motif the measure detects

Known handed tertiary motifs are detected by the tessellation measure, thereby serving as a positive control for the near-achirality found for ordinary packing. The beta-alpha-beta unit, in which two parallel strands are connected through an alpha-helix, is classically of one crossover handedness almost without exception (Richardson, 1976, doi: [10.1073/pnas.73.8.2619](https://doi.org/10.1073/pnas.73.8.2619); Sternberg & Thornton, 1976, doi: [10.1016/0022-2836(76)90099-1](https://doi.org/10.1016/0022-2836(76)90099-1)). I measured a parity-odd crossover descriptor, the signed triple product of the mean strand direction, the strand-to-strand vector, and the offset of the helix from the sheet, over 451 such units, in the non-redundant 846-structure set, in which the strands were parallel and spatially paired. Of these, 98.9% share a single sign (**Fig. S1**), recovering the near-universal right-handed crossover rule, in sharp contrast to the achiral pool of all helix-strand contacts (helix-strand control). The remaining 1.1% carried the opposite handedness, consistent with left-handed crossovers being rare rather than absent. Signed instead by the four-body Delaunay volume of the same crossover tetrahedra, on the enlarged AlphaFold Rossmann-fold set of **Fig. S1**, the motif is only 69% one-handed, so the tessellation measure registers the crossover’s handedness but far more weakly than the directional descriptor.

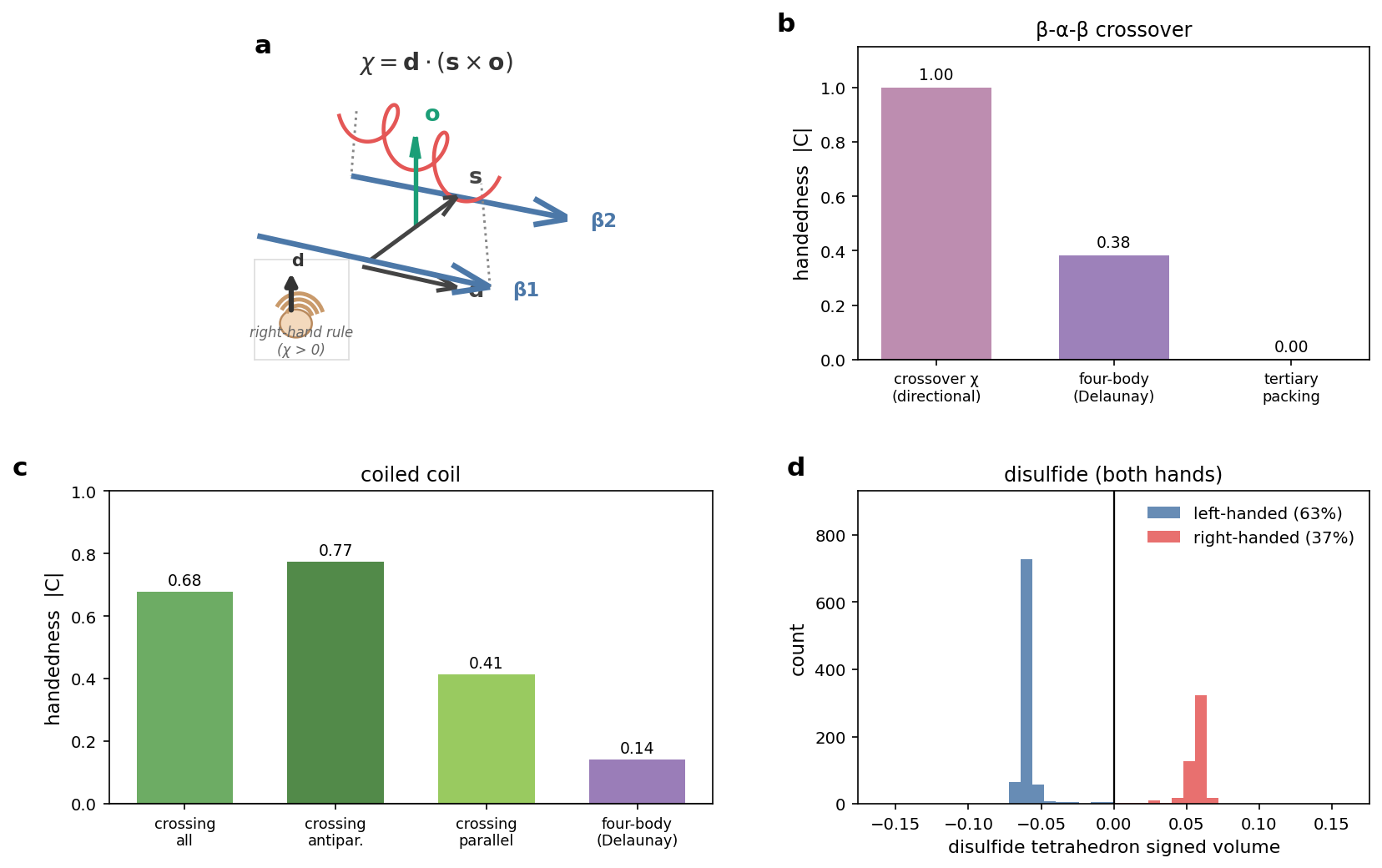

**Figure S1.** Supersecondary-motif handedness, on AlphaFold models. (a) The β-α-β crossover and its directional descriptor *χ* = *d*·(*s*×*o*), with *d* the mean strand direction, *s* the strand-to-strand vector, *o* the helix offset (*χ* > 0 right-handed). (b) β-α-β crossover (Rossmann-enriched set): directional descriptor (|*C*| = 1.00), Delaunay four-body signed volume of the crossover tetrahedra (|*C*| = 0.38), and generic helix-strand packing (|*C*| ≈ 0). (c) Coiled-coil handedness: crossing angle (|*C*| = 0.68) and inter-helix four-body signed volume (|*C*| = 0.14). (d) Disulfide handedness: signed volume of the {Cβ, Sγ, Sγ, Cβ} tetrahedron versus the *χ*3 dihedral (left-handed 63%, right-handed 37%). Descriptors and datasets in Supplementary Methods.

#### Disulfide bonds: individually chiral, balanced as a population

Disulfide bonds give a complementary illustration of the distinction between per-instance chirality and population balance, using a non-tessellation descriptor. The chi3 dihedral Cbeta-Sgamma-Sgamma-Cbeta of a disulfide is bimodal at approximately plus and minus 90 degrees and essentially never near zero, so every disulfide is individually and strongly chiral; the two screw senses are the right-handed (chi3 about +90 degrees, the P helicity) and left-handed (chi3 about -90 degrees, the M helicity) disulfides, both long known to be well represented in the Protein Data Bank (Schmidt, Ho & Hogg, 2006, doi: [10.1021/bi0603064](https://doi.org/10.1021/bi0603064)). Across 1,039 S-S bonds in the non-redundant set of 846 structures the population is only weakly skewed (44.1% right-handed, 95% CI 39.4 to 48.7 on the 846 set, versus 36.6% of 1,382 bonds in the enlarged AlphaFold set of **Fig. S1**), with both handednesses abundantly populated and a slight left-handed excess consistent with the literature in both. Because both signs occur abundantly in natural L-proteins, the disulfides of a reflected D-protein coincide with configurations already present and are in distribution, even though each bond is unambiguously handed. This is the same logic that governs the composition coverage in the main text: what determines whether a mirror configuration is in distribution is population balance, not per-instance chirality.

The β-α-β crossover, coiled-coil, and disulfide handedness controls of **Figure S1** were computed on AlphaFold DB monomer models. β-α-β crossovers were detected as sequential strand-helix-strand element triples with parallel, spatially paired strands, and signed both by the crossover triple product χ = d·(s×o) and by the normalized Delaunay signed volume of the crossover tetrahedra, over 395 Rossmann-fold accessions (UniProt NAD, NADP or FAD keyword) and 487 background human accessions. Coiled coils were 500 UniProt coiled-coil-annotated human accessions; intramolecular helix pairs in sustained close contact with near-parallel or antiparallel axes at coiled-coil spacing were signed by the helix-crossing angle and by the inter-helix four-body signed volume. Disulfides were all cysteine Sγ-Sγ pairs within 2.5 Å across these models; the Cβ-Sγ-Sγ-Cβ tetrahedron signed volume matched the χ3 dihedral in sign for all 1,382 bonds. Handedness is reported as |*C*| = |2f − 1| with f the majority-sign fraction. Detection scripts: delaunay_bab_compute.py, cc_handedness_compute.py, disulfide_compute.py, make_ss_fig.py.

#### Sidechain atomic environments are nearly achiral

To locate the chirality empirically, I began with a purely local descriptor: for each interface atom, the signed triple product of the vectors to its three nearest neighbors (**Fig. S2**). Around side-chain atoms this quantity is symmetric within error (50.0% positive, 95% CI 49.8-50.2; symmetry 0.980), and over all interface atoms it was only marginally skewed (51.7% positive, 95% CI 51.4 to 51.9; symmetry 0.933). A positive control computed on the backbone stereocentre itself, the signed volume of the N, C and Cbeta atoms about the Cα, was by contrast almost entirely one-sided (99.9% positive; symmetry 0.002). The local handedness that a model would encounter is therefore concentrated at the backbone, while the side-chain atomic environment is achiral within error. The side-chain shells of a D-protein thus coincide with shells already present in natural L-structures.

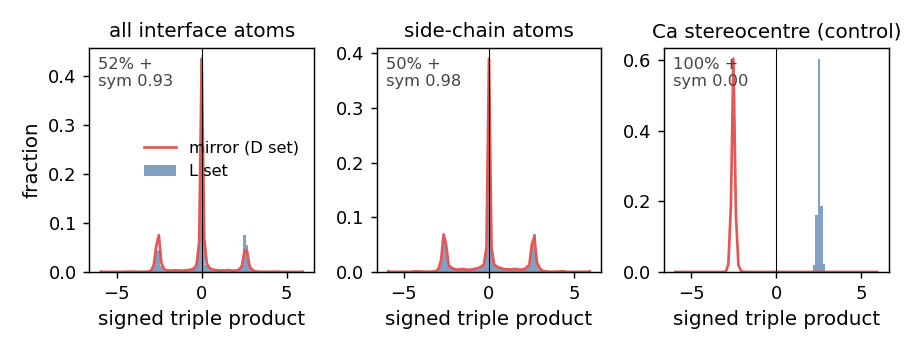

**Figure S2.** Local-shell parity symmetry over the non-redundant set (846 interfaces). Histograms of the signed triple product of the three nearest neighbors of each interface atom (blue), overlaid with the mirror distribution the reflected D-set would produce (red), for all-atom and side-chain environments and, as a control, the Cα stereocentre.

Chirality might nonetheless survive in the relative arrangement of functional groups even where individual atoms appear achiral. To test this possibility, I measured a parity-odd descriptor of the relative orientation of pairs of aromatic side chains on the target surface, constructed from the centroid of each ring and the direction of its Cbeta-to-Cgamma bond so that the internal symmetry of the ring itself does not contribute. The non-redundant set showed no significant handedness (50.8% of one sign, 95% CI 49.3 to 52.2; symmetry 0.932). Side-chain geometry, whether described atom by atom or by the orientation of functional groups, is therefore achiral within error.

#### Composition resolved by vertex order, and robustness to dataset size

A pooled distribution of *V*_D_ can be symmetric either because each composition is two-sided or because one-sided compositions cancel, and the sign of the signed volume requires an ordering of the four vertices, so the near-symmetry could in principle reflect that labeling. Both possibilities are tested here by binning every tertiary tetrahedron by its unordered composition and, separately, by its ordered composition (the residue at each vertex in sequence order i < j < k < l), across the non-redundant set of 846 complexes and the larger ProteinMPNN split of about 20,000 structures (Methods).

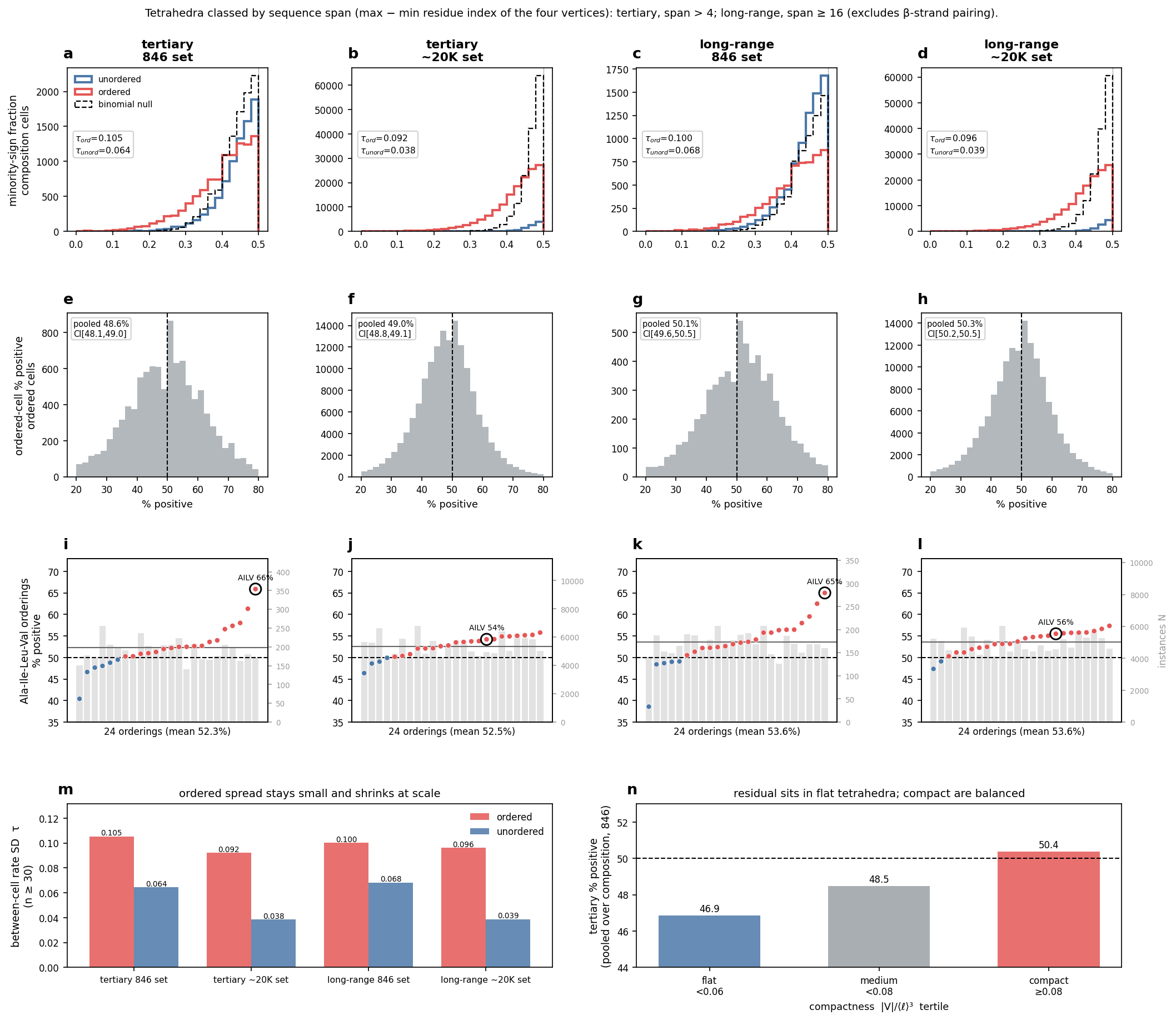

**Figure S3.** Per-composition two-sidedness is robust to vertex ordering, span, and dataset size. Columns are four (span stratum, dataset) combinations: tertiary/846, tertiary/~20K, long-range/846, long-range/~20K (tertiary, span > 4; long-range, span ≥ 16; cells with ≥ 30 tetrahedra). (a-d) Per-cell minority-sign fraction min(+,−)/total for unordered (blue) and ordered (red) composition, with a matched binomial null (dashed); between-cell rate SD *τ* annotated. (e-h) Per-cell % positive across ordered cells (dashed line at 50%); inset, pooled positive fraction with 95% structure-bootstrap CI. (i-l) The 24 sequence orderings of Ala-Ile-Leu-Val sorted by % positive (points; grey bars, instance count *N*), canonical ordering circled. (m) Between-cell rate SD *τ* for ordered versus unordered cells across the four combinations. (n) Tertiary % positive stratified by compactness |*V*|/⟨ℓ⟩³ into flat, medium, and compact tertiles (846 set). Combination counts in Table S1.

***Table S1.*** *Number of composition cells populated and analyzed in the* ***Fig. S3*** *control, for each span stratum and dataset. Cells are single compositions taken either as an unordered multiset of four residue types or as an ordered tuple (a residue at each sequence-order slot i < j < k < l). With 20 amino acid types the ceilings are 8,855 possible unordered compositions and 20^4^ = 160,000 possible ordered compositions; the n >= 30 columns count the cells retained for the per-cell statistics (at least thirty tetrahedra). The final column gives the total number of tertiary or long-range tetrahedra. The ordered space is nearly saturated on the ~20K set but sparsely sampled on the 846 set, which is why the ordered Ala-Ile-Leu-Val cell in* ***Fig. S3*** *regresses from 66% to 54% with scale.*

| **Combination** | **Ordered cells (populated)** | **Ordered cells (n >= 30)** | **Unordered cells (populated)** | **Unordered cells (n >= 30)** | **Tetrahedra** |
| --- | --- | --- | --- | --- | --- |
| tertiary / 846 | 146,804 | 10,270 | 8,852 | 8,070 | 1,740,985 |
| tertiary / ~20K | 159,995 | 152,429 | 8,855 | 8,854 | 47,394,354 |
| long-range / 846 | 141,502 | 6,474 | 8,850 | 7,729 | 1,382,889 |
| long-range / ~20K | 159,989 | 147,979 | 8,855 | 8,854 | 38,120,102 |

#### Vertex ordering: how the achiral orderings differ

The sign of a tetrahedron's signed volume depends on the order of its four vertices, so the near-achirality could in principle depend on the ordering rule. Three sign-sensitive orderings are compared here: the sequence order used in the main analyses, and two shape-intrinsic orderings (by distance to the tetrahedron centroid and by summed distance to the other three vertices) that never use the sequence.

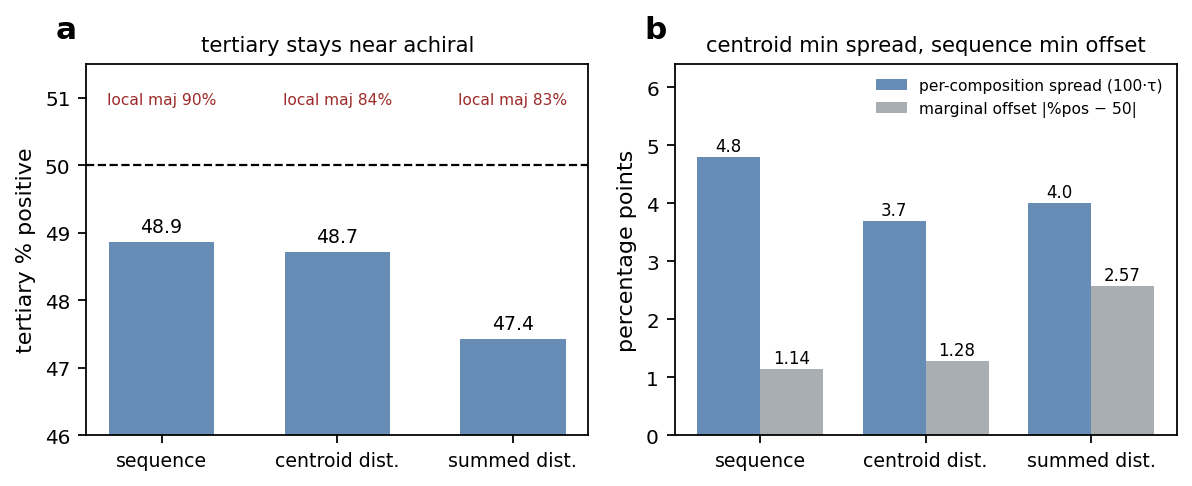

**Figure S4.** How the three sign-sensitive vertex orderings differ for tertiary packing (per-composition spread and marginal from the 20,383-structure split; local marginals from the 846 set). (a) Tertiary % positive under each ordering, all within one to three percent of 50; the sequence-local majority-sign percentage is annotated for each, confirming every ordering is strongly one-sided locally (the sign is a convention). (b) The between-composition rate spread *τ* and the marginal offset |dev| for each ordering.

#### Fold handedness is widespread but cancels across the ensemble

Individual protein folds are frequently handed at the tertiary scale, yet the ensemble of folds cancels to near-zero (**Fig. 5**). A random sample of X-ray structures was domain-isolated to its largest CATH domain and scored for tertiary *C* over tetrahedra of sequence span ≥ 8 (823 scorable structures, 272 CATH superfamilies, 27 architectures); domain isolation removes globular parts of multidomain chains that would otherwise dilute the signal. Figure S5 ranks 30 common folds (CATH superfamilies sampled fresh from the PDB), each with a 95% structure-bootstrap confidence interval (3,000 replicates). Code and identifier lists are in the reproducibility package.

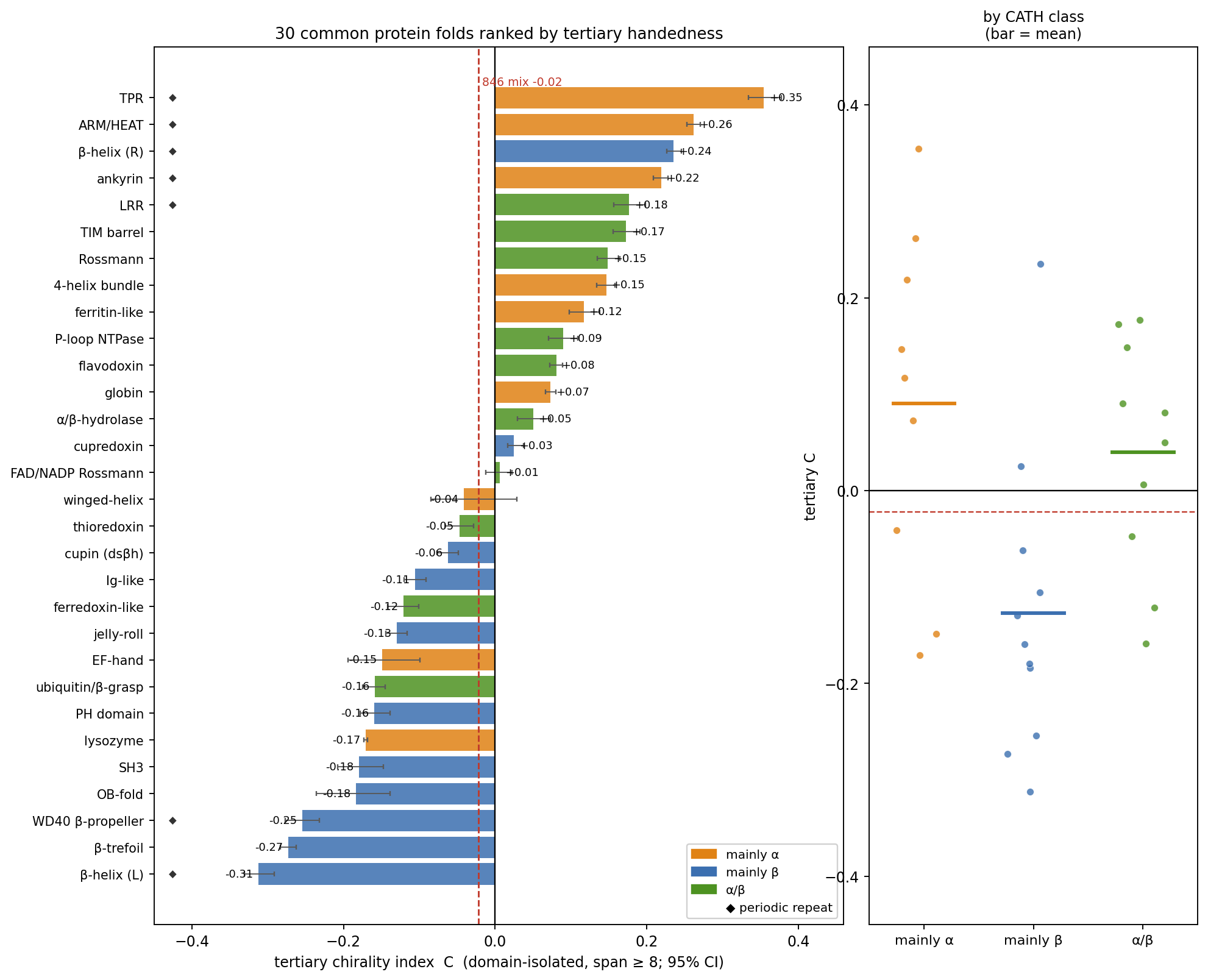

**Figure S5.** Tertiary handedness ranked across 30 common protein folds. Each fold is a CATH superfamily sampled fresh from the PDB and domain-isolated; bars are the pooled domain-isolated tertiary chirality index *C* (span ≥ 8) with a 95% structure-bootstrap confidence interval (3,000 replicates), colored by CATH class. Solenoids and repeat folds occupy the extremes, β-sheet folds skew negative and α and α/β folds positive, and the set straddles zero.

*Table S2. Reproducibility map. Each display item is regenerated by running its compute script(s) on the listed id list (python3 scripts/<compute>.py --idlist <file> --out data/<name>.jsonl, called repeatedly until complete), then its plotting script (python3 scripts/<plot>.py). All compute scripts share the same --idlist / --out / --max-seconds interface. The non-redundant PPI set is the mpnn_ppi_ids.txt list of 846 complexes; Figure 4 also uses the ProteinMPNN composition set of 20,383 structures (mpnn_ids.txt) and Figure 3 the LigandMPNN small-molecule set.*

| **Item** | **Compute script(s) (--idlist)** | **Id-list file** | **Plotting script** |
| --- | --- | --- | --- |
| Fig 1 | make_schematic.py (illustrative, PDB 1YCR) | none | make_schematic.py |
| Fig 2 | delaunay_inter_compute.py, delaunay_span_compute.py; make_idealized_ss.py (panel a idealized models and signed-volume histograms) | mpnn_ppi_ids.txt | make_ss_fig.py |
| Fig 3 / Table 1 | delaunay_nr_compute.py, delaunay_env_compute.py, ligand_site_run.py, delaunay_span_compute.py (datasets: also nr_ids.500.txt) | mpnn_ppi_ids.txt (ligand: ligand_small_ids.txt) | make_headline_fig.py |
| Fig 4 | delaunay_repr_compute.py, delaunay_ordering_compute.py, mpnn_comp_run.py, comp_shape_compute.py | mpnn_ppi_ids.txt (comp: mpnn_ids.txt) | make_repr_fig.py |
| Fig 5 | random_pdb_fold_scan.py (CATH-domain isolated) | random PDB sample (+ CATH domains) | fig5 histogram; pymol_fig5_panels.py |
| Fig 6 | make_tradeoff_schematic.py (schematic; y-axis anchored by measured \|*C*\|) | none | make_tradeoff_schematic.py |
| Fig S1 | delaunay_bab_compute.py (helix-strand control: delaunay_inter_compute.py) | mpnn_ppi_ids.txt | make_ss_fig.py |
| Fig S1 | disulfide_compute.py | mpnn_ppi_ids.txt | disulfide_aggregate.py |
| Fig S2 | nr_lenses_compute.py | mpnn_ppi_ids.txt | nr_lenses_aggregate.py |
| Fig S3 | comp_ordered_control.py, comp_ordered_stats.py, comp_shape_ordered.py | mpnn_ppi_ids.txt (scale: mpnn_ids.txt) | make_ordered_control_fig.py |
| Fig S4 | comp_ordering_run.py | mpnn_ids.txt | make_ordering_si_fig.py |
| Fig S5 | scan_cath_folds.py; make_fold_ranking.py | 30 CATH superfamilies | make_fold_ranking.py |
| Fig S6 | architecture_census.py | CATH architectures | architecture_census.py |
